# CRISPR-based editing of the ω- and γ-gliadin gene clusters reduces wheat immunoreactivity without affecting grain protein quality

**DOI:** 10.1101/2023.01.30.526376

**Authors:** Zitong Yu, Ural Yunusbaev, Allan Fritz, Michael Tilley, Alina Akhunova, Harold Trick, Eduard Akhunov

## Abstract

Wheat immunotoxicity is associated with abnormal reaction to gluten-derived peptides. Attempts to reduce immunotoxicity using breeding and biotechnology often affect dough quality. Here, the multiplexed CRISPR-Cas9 editing of cultivar Fielder was used to modify the ω -and γ-gliadin gene clusters abundant in immunoreactive peptides identified in the genomes assembled using the long-read sequencing technologies. The whole genome sequencing of an edited line showed editing or deletion of nearly all ω-gliadin and half of the γ-gliadin gene copies and lack of editing in the α/β-gliadin genes. The detected 62% and 52% reduction in ω- and γ-gliadin content, respectively, had no negative impact on grain protein quality. A 47-fold immunoreactivity reduction compared to wild-type was detected using antibodies against immunotoxic peptides. Our results indicate that genome profiling to identify gliadin gene copies abundant in immunoreactive peptides and their targeted editing could be an effective mean for reducing immunotoxicity of wheat cultivars while minimizing the impact of editing on protein quality.

## Introduction

Wheat gluten defines the breadmaking properties of dough and underlies immune reaction to wheat-based products (Shewry and Tatham, 2016). The efforts to reduce wheat immunotoxicity often negatively affected dough quality (Gil-Humanes et al., 2010; 2014). The question remains whether it is feasible to selectively modify copies or parts of gluten-encoding genes to substantially reduce wheat immunotoxicity without affecting the physicochemical properties of dough important for breadmaking.

The gluten network/gluten macropolymer (GMP) formed by disulphide bonds (DBs) between the cysteine residues of glutenins and gliadins defines the quality of dough (Shewry and Tatham, 1997). The inter-molecular DBs between high molecular weight glutenin subunits (HMW-GS) extend gluten network, whereas intra-molecular DBs formed by gliadins terminate glutenin polymerization (Shewry and Halford, 2002). The impact of ω-gliadins, which lack cysteine residues, on the gluten network is expected to be lower than that of α/β- and γ-gliadins, which could form three and four intramolecular DBs, respectively (Visschers and de Jongh, 2005; Wieser, 2007). This is consistent with improved dough mixing time and tolerance observed in the RNAi transgenic lines with the ω1,2-gliadin expression suppressed (Altenbach et al., 2019). Considering that ω-gliadins represent only a small fraction of all gliadins, the CRISPR-Cas9 editing of these genes will likely have limited negative impact on dough quality.

Celiac disease, food allergy and wheat-dependent exercise-induced anaphylaxis (WDEIA) are associated with reaction to gluten induced by the oral intake of wheat gluten components resistant to enzymatic degradation. As a result, the repetitive peptides rich in proline (Pro, P) and glutamine (Gln, Q) accumulate in the small intestine (Scherf et al., 2016). Among gluten fractions, gliadins are the major carriers of toxic epitopes. The genes encoding ω- and γ-gliadins are tightly clustered at three loci *Gli-A1*, *Gli-B1* and *Gli-D1* on chromosomes 1A, 1B and 1D, respectively. The genes encoding α/β-gliadins are located on the short arm of chromosome 6, named *Gli-A2*, *Gli-B2* and *Gli-D2* (Branlard et al., 2001). Celiac disease and food allergy are caused by all three gliadin subtypes; while the WDEIA is associated with ω5-gliadins encoded by the genes located at *Gli-B1* loci (Shewry and Tatham, 2016). The ω1,2-gliadins encoded by the genes located at *Gli-A1* and *Gli-D1* loci contain multiple peptides enriched in repeats containing the highly immunoreactive QQPFP motif. Food allergy-related toxic epitopes are prominent in all gliadin subtypes, with the highest number present in ω-gliadins, followed by γ- and α/β-gliadins. Peptides including PQQPFP, QPQQPFP, and QQFPQQQ motives are frequently linked with food allergy. The toxic epitopes including QQFPQQQ motif associated with WDEIA are most frequent in ω5-gliadins located on chromosome 1B (Juhász et al., 2018).

A reduction of gluten toxicity is often accompanied by decrease in gluten quality. The RNAi lines showing 80% reduction in γ-gliadin expression and reduced reaction against the R5 monoclonal antibody (mAb) also showed reduced dough strength (Gil-Humanes et al., 2008; 2012). Likewise, 62-67% reduction in immunoreactivity to the R5 and G12 mAbs achieved by the CRISPR-Cas9 editing of 35 of 45 α-gliadin genes was accompanied by decrease in SDS sedimentation volume and lower dough quality (Sánchez-León et al., 2018). However, the RNAi suppression of ω1,2-gliadins significantly reduced reactivity of flour proteins to serum IgG and IgA antibodies and improved both mixing time and tolerance in transgenic lines (Altenbach et al., 2019). This result suggests that removal of certain epitopes remains feasible without detrimental impact on protein quality.

Accurate annotation of multigene families in complex wheat genome is critical for their successful editing. The recently released genome assembly of the transformation-amenable cultivar Fielder (Sato et al., 2021) generated using HiFi reads provide unique opportunity for identifying gliadin genes enriched for immunoreactive peptides and designing guides for their targeted editing. Here, we analyzed the distribution of 11 immunoreactive peptides binding to R5 mAb and six peptides binding to G12 mAb (Schopf and Scherf, 2018) across the three gliadin gene families annotated in four high-quality genome assemblies of cultivars Fielder (Sato et al., 2021), Kariega (Athiyannan et al., 2022), Chinese Spring and LongReach Lancer (Walkowiak et al., 2020). Since most ω-gliadins lack cysteine residues to form disulphide crosslinks in gluten network, and the proportion of ω- and γ-gliadins in gluten is lower than that of α/β-gliadins, we expect that CRISPR-Cas9-mediated editing of ω- and γ-gliadin gene clusters should reduce gliadin toxicity and maintain protein quality. The multiplex editing was performed in cv. Fielder using gRNAs targeting ω- and γ-gliadin gene clusters. The whole genome sequencing of an edited line revealed large- and small-scale deletions in gliadin gene clusters or individual genes. The analyses of gliadin protein content and SDS-extractable and -unextractable polymeric protein profiles demonstrated significant reduction in ω- and γ-gliadin protein content and lack of impact on protein quality. The monoclonal antibodies showed 47-fold reduction in immunoreactivity of gluten in edited lines compared to wild-type Fielder. These results indicate that improved genome assemblies and annotation make feasible targeted editing of complex gene cluster in wheat and provide opportunities for reducing its immunotoxicity without affecting end-use quality traits.

## Results

### Distribution of immunogenic peptides across gliadin genes in cv. Fielder genome

To investigate distribution of peptides associated with immune response across gliadin gene clusters, we have annotated the cv. Fielder genome (see Materials and Methods). We have identified five, eight and five copies of ω-gliadin genes in chromosomes 1A, 1B and 1D, respectively. Translation of these ω-gliadin genes suggest that two, two and five copies on 1A, 1B and 1D, respectively, are functional. Out of seven, eight and nine copies of the γ-gliadin genes in chromosomes 1A, 1B and 1D, respectively, five, seven and eight copies had uninterrupted open reading frame. For α/β-gliadin genes, there are 11, 27 and 14 copies on chromosomes 6A, 6B and 6D, respectively, with 10, 15 and nine copies being functional.

The R5 mAb raised against rye secalin binds to 11 gliadin toxic epitopes including QQPFP, QLPFP, LQPFP, QLPYP, QLPTF, QQSFP, QQTFP, PQPFP, QQPYP, QQQFP, QVQWP; the G12 mAb raised against the 33-mer gliadin peptide can interact with six toxic epitopes including QPQLPY, QPQLPF, QPQLPL, QPQQPY, QPQQPF, QPELPY (Schopf and Scherf, 2018; Fig. 1a). We analyzed the distribution of 11 immunoreactive peptides binding to R5 mAb and six peptides binding to G12 mAb across the three gliadin gene families annotated in four high-quality assemblies of cultivars Fielder (Sato et al., 2021), Kariega (Athiyannan et al., 2022), Chinese Spring and LongReach Lancer (Walkowiak et al., 2020) (Fig. 1a; Supplementary Figure 1). The mapping result shows that the number of R5- and G12-binding peptides in ω- / γ-gliadins was 1.1 – 1.8 and 4.7 – 9.8 times higher, respectively, than that in α/β-gliadins, with 487 vs 263 and 220 vs 26 in Fielder, 358 vs 245 and 136 vs 22 in Kariega, 194 vs 185 and 84 vs 18 in Chinese Spring, and 198 vs 156 and 78 vs 8 in LongReach Lancer (Fig. 1b; Supplementary Table 1). These results suggest that substantial reduction in the R5- and G12-based immunoreactivity could likely be achieved by editing the ω- and γ-gliadin genes.

**Figure 1.**
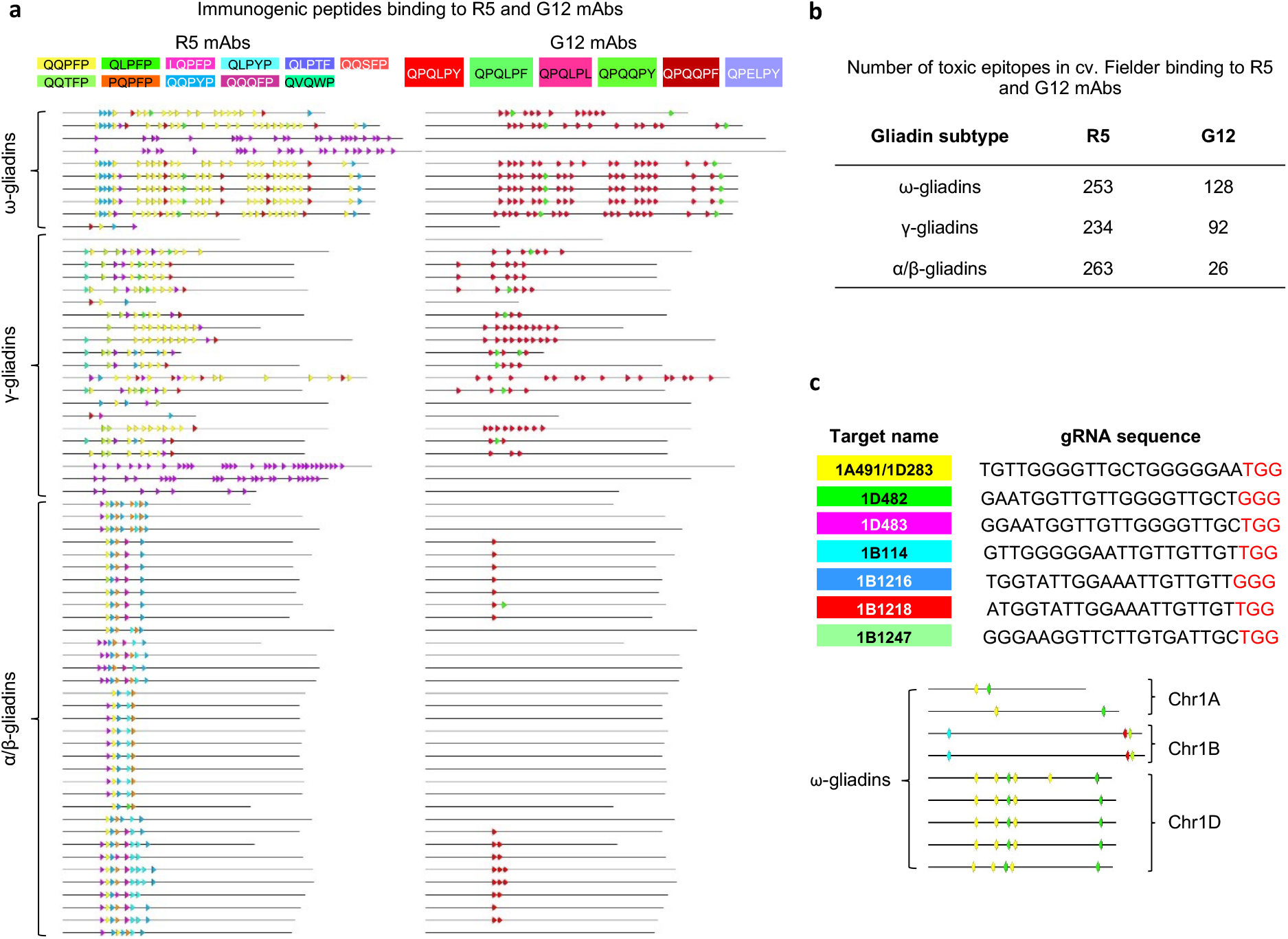
Distribution of toxic epitopes binding to R5 and G12 mAbs across gliadin genes in cultivar Fielder. **a**. The amino acid motifs of the 11 toxic epitopes binding to R5 mAb and the six toxic epitopes binding to G12 mAb: the 17 motifs were indicated by different background colors. The distributions of the 11 toxic epitopes binding to R5 mAb (left) and the six toxic epitopes binding to G12 mAb (right) across all three gliadin subtypes of cultivar Fielder: the different toxic epitopes were indicated by the arrows with corresponding color shown in Fig. 1a; the distributions of the 17 toxic epitopes binding to R5 and G12 mAbs across all three gliadin subtypes of cultivars Kariega, Chinese Spring and LongReach Lancer were shown in Supplementary Fig. 1. **b**. The number of toxic epitopes binding to R5 and G12 mAbs in each gliadin subtype of cultivar Fielder; the number of peptides binding to R5 and G12 mAbs from all gliadin subtypes of cultivars Kariega, Chinese Spring and LongReach Lancer is shown in Supplementary Table 1. **c**. Seven gRNAs designed for gliadin gene editing: three gRNAs target ω-gliadins on chromosomes 1A and 1D and four gRNAs target ω-gliadins on chromosome 1B. The PAM site NGG was indicated by the red font; the distribution of seven gRNAs across nine functional copies of the ω-gliadin genes including two on chromosome 1A, two on chromosome 1B and five on chromosome 1D. The color coding for gRNA target sites corresponds to colors shown in Fig. 1c. The gRNA targeted regions within the ω- and γ-gliadin genes with 0 to 3 mismatches are shown in Supplementary Table 2.

### CRISPR-Cas9-mediated editing of the ω- and γ-gliadin genes in cv. Fielder

Due to higher number of R5- and G12-binding peptides in ω-gliadins compared to γ- gliadins, the design of guide RNAs (gRNAs) was primarily focused on ω-gliadin genes. Among the ω-gliadin gRNAs, the preference was given to those that can also target some of the γ-gliadin gene copies. A total of three gRNAs were designed to target ω-gliadin loci on chromosomes 1A and 1D, including gRNAs 1A491/1D283, 1D482 and 1D483; and four gRNAs were designed to target loci on chromosome 1B, including gRNAs 1B114, 1B1216, 1B1218 and 1B1247 (Fig. 1c). These gRNAs had from 0 to 3 mismatches in the targeted repetitive regions within the ω- and γ- gliadin genes (Supplementary Table 2). Each gRNA was subcloned into a separate Csy4-spCas9 vector, combined in equimolar proportions into single pool and co-transformed into cv. Fielder.

A total of 435 T_1_ plants were obtained from 32 tillers of transgenic Cas9-positive T_0_ plants. Three PCR primer pairs were used to screen plants for the presence of gene edits in the nine functional copies of the ω-gliadin genes (Supplementary Tables 3 and 4). Due to the complexity and high levels of similarity, we experienced difficulties with designing specific primers for the γ-gliadin gene copies and opted to use whole-genome sequencing to characterize gene edits in this group of genes for a targeted line (see whole-genome sequencing results below). PCR screening detected fragment deletions in the ω-gliadin genes located on chromosomes 1A and 1D of seven T_1_ plants derived from T_0_ line 387-3, including plants 387-3-1, 387-3-2, 387-3-3, 387-3-4, 387-3-5, 387-3-6, and 387-3-7 (Supplementary Fig. 2), while no fragment deletions were found in the gene copies on chromosome 1B. The next generation sequencing (NGS) showed that T_0_ line 387-3 was transformed with four gRNAs, including 1A491/1D283, 1B114, 1B1247, and 1D482. The T_1_ progenies of 387-3 was negative for Cas9 sequence, except for one plant (387-3-7). The PCR analysis of the T_2_ population including 183 lines derived from plant 387-3-6 confirmed the absence of segregation for the fragment deletions (Supplementary Fig. 3), revealing that the T_1_ line 387-3-6 is homozygous for the detected gene edits. The analyses of editing events were conducted in the T_1_ line 387-3-6.

The initial characterization of editing events in line 387-3-6 was performed by the NGS of PCR amplicons generated for the ω-gliadin genes. The fragment deletions ranging from 18 to 111-bp were detected in one, two and five copies of ω-gliadin genes on chromosomes 1A, 1B and 1D, respectively (Supplementary Fig. 4; Supplementary Table 5). Nearly all reads mapping to these gliadin loci carried mutated variants confirming that line 387-3-6 is homozygous for the edited alleles of the ω-gliadin genes. Because designing PCR primers for amplifying all gene copies in the ω- and γ-gliadin gene clusters was not feasible due to their repetitive nature, more detailed analysis of on- and off-target editing events in 387-3-6 line was conducted by whole genome sequencing.

TruSeq DNA PCR-free library prepared for line 387-3-6 was sequenced at 10x genome coverage using Illumina technology (2 x 150-bp). First, by mapping reads to the Cas9-gRNA constructs we confirmed that line 387-3-6 is free of the transgenic sequences used for wheat transformation. Then, the reads were mapped to the recently released genome of cultivar Fielder generated using the long-read PacBio sequencing data (Sato et al., 2021). The genome annotation identified nine and 22 functional ω- and γ-gliadin gene copies, respectively. Reads mapping to both unique and multiple locations in genome were used to identify edited ω- and γ-gliadin genes (Supplementary Table 6) based on the changes in the depth of read coverage, gaps in the alignments of assembled gliadin genes, and presence of short indels at the gRNA binding sites (see Methods). We detected two large-scale deletions on chromosomes 1A and 1D that resulted in the loss of two and four functional ω-gliadin genes (Fig. 2), respectively, indicating that targeted removal of gliadin gene clusters is feasible in wheat using the gene editing approach. The smaller scale deletions affecting either portion of the genes or regions at the gRNA binding sites were detected in remaining three functional copies of the ω-gliadin genes and 12 functional copies of the γ-gliadin genes (Fig. 2). These results are consistent with the observed reduction of ω- and γ-gliadin content in flour protein extracts (Fig. 3b). The loss of gliadin genes due to editing resulted in removal of 55.2% and 77.2% of immunotoxic peptides binding to R5 and G12 mAbs identified in the Fielder genome (Fig. 3e). The analysis of potential off-target editing sites that have up to three mismatches with the designed gRNAs revealed lack of Cas9-induced mutations outside of the gliadin genes, indicating that the developed line does not carry undesirable modifications in the genome.

**Figure 2.**
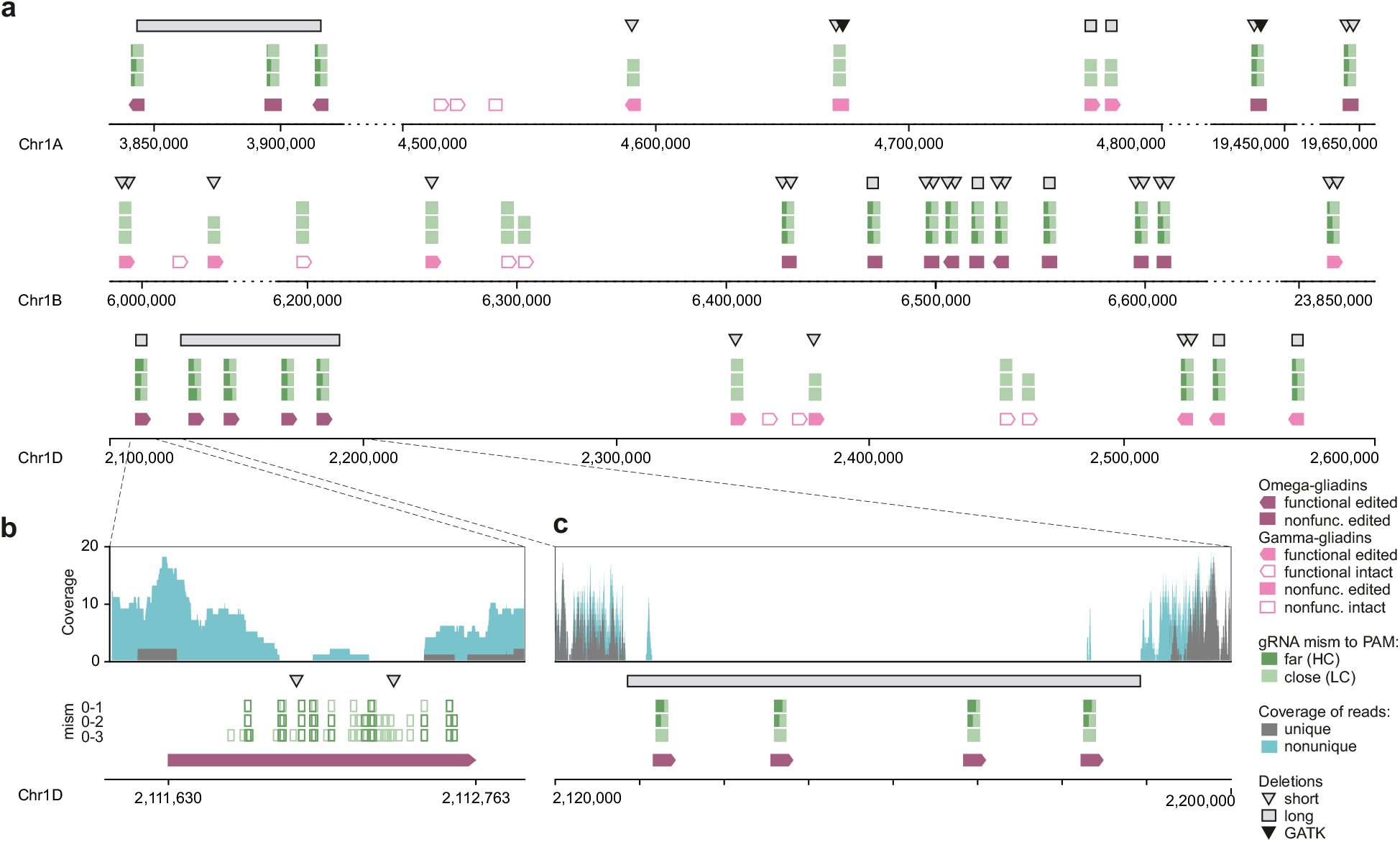
Summary of gene editing events identified by whole genome sequencing in the ω- and γ-gliadin gene clusters. **a**. Distribution of gRNA target sites and different types of gene editing events among the ω- and γ-gliadin genes on chromosomes 1A, 1B and 1D. **b, c**. The distribution of the depth of read coverage, the types of editing events, and gRNA target sites with 0, 1, 2 or 3 mismatches to the gRNA protospacers for the ω-gliadin genes with the partial deletion of coding region (b) and deletion of a cluster of four genes (c).

**Figure 3.**
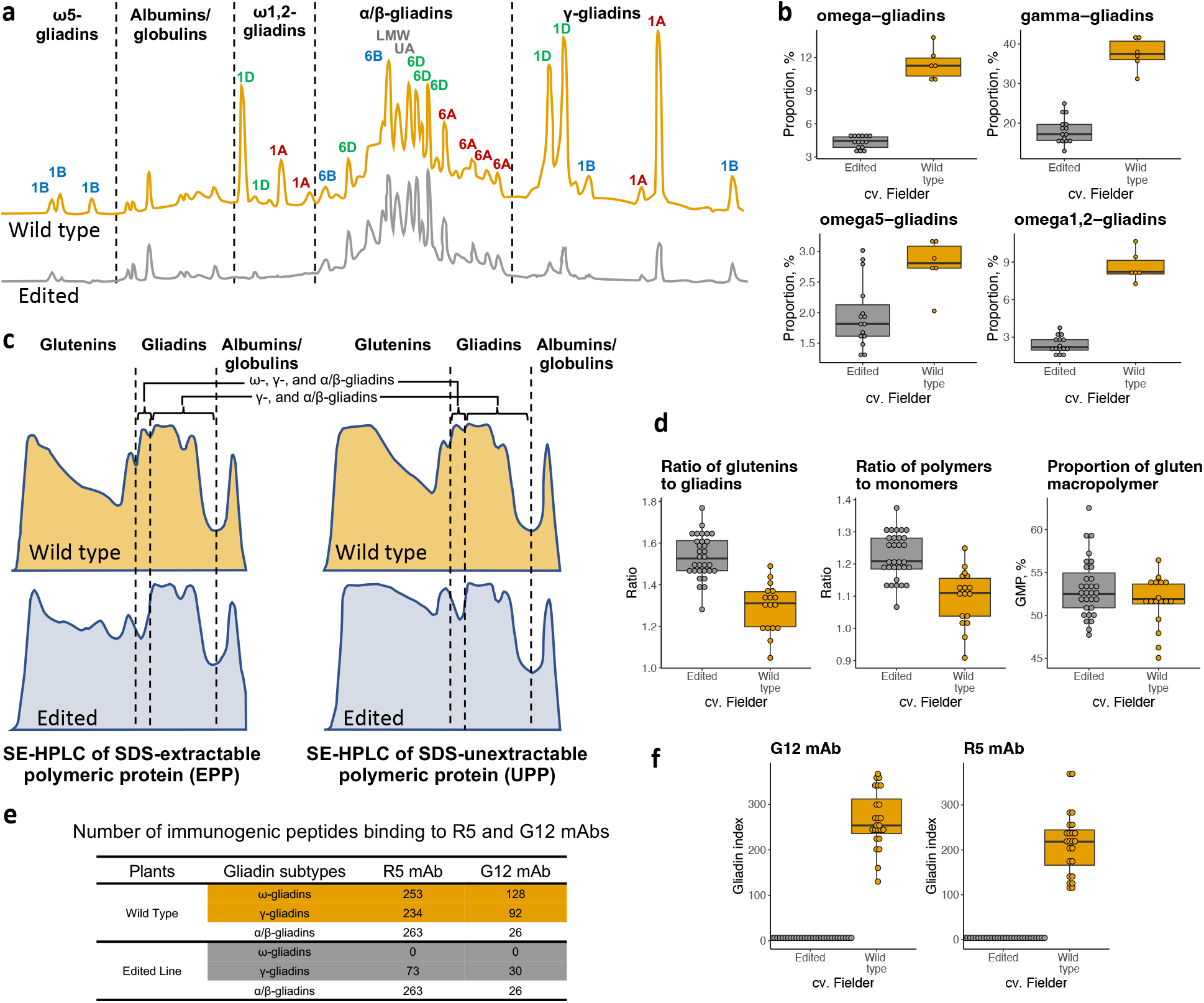
The impact of gene editing on gliadin content, properties of grain protein and immunoreactivity. **a**. RP-HPLC based gliadin profiles of wild-type Fielder and a T_2_ plant derived from the edited line 387-3-6. Based on the hydrophobicity of each component (five areas separated by the black dotted lines) were assigned to ω5-gliadins, albumins/globulins, ω1,2-gliadins, α/β-gliadins and γ-gliadins. The peaks encoded by the genes located on chromosomes 1A, 1B and 1D are labeled using red, blue and green fonts. **b**. Boxplots show the proportion of ω- and γ-gliadin subtypes in the gliadin extracts from wild-type cv. Fielder and edited lines. The individual datapoints for six wild-type samples and 15 randomly selected edited lines are shown. **c**. SE-HPLC based SDS-extractable (left) and SDS-unextractable (right) polymeric protein profiles of wild-type Fielder and two randomly selected T_2_ plants derived from edited line 387-3-6. Based on the molecular weight of each component, three areas separated by black dotted lines were assigned to glutenins, gliadins and albumins/globulins. **d**. Comparisons of glu/gli ratio, poly/mono ratio and GMP % between the wild-type Fielder and the edited lines. The datapoints are shown for 17 wild-type samples and 62 randomly selected edited line derived from T_1_ plant 387-3-6. **e**. Comparison of the number of immunogenic peptides binding to the R5 and G12 mAbs in wild-type Fielder and edited lines. The number of toxic epitopes in the gene edited line is estimated using the intact coding regions of gliadin genes identified by whole genome sequencing. **f**.Comparison of immunoreactivity between the non-edited and edited lines measured using R5 and G12 mAbs. The datapoints are shown for 24 non-edited lines and 30 randomly selected edited lines derived from T_1_ plant 387-3-6. Gliadin index corresponds to gliadins × 10^4^ ppm. In the figure, boxplots show the mean value and interquartile ranges (IQR); the line inside the box indicated the mean value; the end of the top line was the third quartile (Q3) + 1.5× IQR; the end of the bottom line was the first quartile (Q1) – 1.5× IQR.

### The impact of gene editing on gliadin content and properties of grain protein

Reverse-phase high performance liquid chromatography (RP-HPLC) technology was used to compare the amount of ω- and γ-gliadins in the T_2_ plants of edited line 387-3-6 with wild-type Fielder (Fig. 3a). The analysis revealed statistically significant decrease of ω- and γ-gliadins in the gene edited line. The total amount of ω-gliadins was decreased by 61.9% (*t*-test; p-value = 3.59E-05; Fig. 3b), with ω1,2-gliadins and ω5-gliadins decreased by 72.3% and 29.5%, respectively (*t*-test; p-value = 1.47E-05 and 3.00E-03; Fig. 3b). The amount of γ-gliadins was decreased by 51.7% (*t*-test; p-value = 4.95E-06; Fig. 3b). So, here we confirm that the amount of ω- and γ-gliadins are substantially reduced by CRISPR-Cas9 mediated gene editing.

To assess the impact of gene editing on breadmaking properties of dough, we have analyzed the SDS-extractable polymeric proteins (EPP) and SDS-unextractable polymeric proteins (UPP) using size-exclusion HPLC (SE-HPLC) (Ohm et al., 2009a; 2009b). The SE-HPLC data was used to calculate three parameters of protein extracts that strongly correlate with breadmaking quality: the ratio of glutenins to gliadins (glu/gli), the ratio of polymeric to monomeric proteins (poly/mono) and the percentage of GMP (GMP %). The first peak corresponding to gliadins in the profiles of EPP and UPP was missing in the gene edited line compared to wild-type (Fig. 3c). The glu/gli and poly/mono ratios increased from 1.29 to 1.53 (*t*-test; p-value = 6.10E-08; Fig. 3d) and from 1.10 to 1.22 (*t*-test; p-value = 1.82E-05; Fig. 3d), respectively, in the gene edited line compared to wild-type. There was slight increase in GMP % from 51.6% to 53.2%, though difference was not statistically significant (*t*-test; p-value = 9.56E-02; Fig. 3d). These results suggest that decrease in the ω- and γ-gliadin content in the gene edited line facilitates gluten polymerization and improves the parameters of flour protein extracts correlating with breadmaking quality.

### The comparison of immunoreactivity between the non-edited and edited lines

The mAbs R5 and G12 raised against secalins and gliadins are highly predictive of wheat immunotoxicity for gluten sensitive patients (Morón et al., 2008, Valdés et al., 2003). We have used 24 Cas9-negative non-edited transgenic lines and 30 edited lines derived from line 387-3-6 to assess the immunoreactivity of gliadin extracts against both the R5 and G12 mAbs. There was 47-fold reduction (97.9%) in the average immunoreactivity of gliadins against R5 mAb in the edited lines (4.58 × 10^4^ ppm) compared to that in the controls (214.83 × 10^4^ ppm) (*t*-test; p-value = 3.44E-13; Fig. 3e, 3f). The immunoreactivity against G12 mAb was reduced from 266.65 × 10^4^ ppm in control lines to 6.29 × 10^4^ ppm in the gene edited lines, which corresponds to 42-fold reduction (97.6%) (*t*-test; p-value =3.68E-16; Fig. 3e, 3f). So, here the results show that the immunoreactivity of gene edited line has a substantial reduction around 97% at up to 42-fold compared to wild-type. By using Total Gluten assay R7041 (Lacorn et al., 2019), which can detect nearly all gluten proteins in wheat (gliadins, HMW- and LMW-GS), we detected approximately 30% immunoreactivity reduction in the edited lines (*t*-test; p-value = 9.81E-7) compared to wild-type Fielder. This result suggests that the edited immunotoxic peptides from the gliadin genes in line 387-3-6 account for about third of total gluten immunoreactivity detectable by the R7041 assay.

## Discussion

Due to the complexity of wheat genome, targeted editing of multigene clusters that might show significant structural differences from the available reference genomes represents a significant challenge. Our study demonstrates that new genomic resources could be combined with a multiplex editing approach for engineering multigene clusters in the wheat genome. The high quality of wheat genome assemblies, especially the genome of the transformation amenable cv. Fielder used in our study for transformation, generated using long-read sequencing technologies (Walkowiak et al., 2020; Sato et al., 2021; Zhu et al., 2021; Athiyannan et al., 2022) provided accurate annotation of the highly repetitive gliadin gene families. This permitted us to investigate the distribution of immunotoxic peptide motifs across all three gliadin subtypes in the genome of cultivar Fielder and design gRNAs for targeted deletion of highly immunoreactive ω- and γ-gliadin gene copies.

The modification of targeted genes was confirmed by whole genome sequencing of the edited wheat line, which detected both small- and large-scale deletions in the ω-and γ-gliadin gene clusters. The two large-scale deletions on chromosomes 1A and 1D led to the loss of two and four functional ω-gliadin genes, respectively. This result indicates that targeted removal of large multigene clusters is feasible in wheat using the gene editing approach (Jouanin et al., 2020), opening opportunities for engineering gluten gene loci and their removal and replacement with either natural or synthetic variants of genes exhibiting lower immunoreactivity and improved quality.

In the previous study, it was shown that CRISPR-Cas9 editing of cv. Bobwhite preferentially targeting α/β-gliadin genes results in increased proportion of glutenin relative to gliadins (Sánchez-León et al., 2018). However, this was accompanied by significant reduction in the amount of GMP measured using SDS sedimentation approach (Sánchez-León et al., 2018), suggestive of reduced breadmaking quality. It is likely that the copies of the α/β-gliadin genes edited in that study played important role in forming the gluten network. Contrary to results obtained for lines with the edited α/β-gliadin genes, the percentage of GMP, which positively correlates with breadmaking quality, was slightly increased in edited line 387-3-6. Substantial increase in the ratio of polymer to monomer in line 387-3-6 also indicates that the deletion of ω- and γ-gliadin gene copies likely had minor negative impact on gluten polymerization. These results suggest that targeted editing of the ω-and γ-gliadin genes in cv. Fielder had positive impact on properties of wheat gluten affecting breadmaking quality of wheat flour.

The R5 and G12 mAbs raised against immunoreactive peptides from secalins and gliadins are predictive of wheat immunotoxicity for gluten sensitive patients (Morón et al., 2008, Valdés et al., 2003). The gluten immunoreactivity to G12 and R5 mAbs in our gene edited lines was similar to that previously reported for gene-edited cultivar Bobwhite (Sánchez-León et al., 2018). However, the reduction of immunoreactivity in our study compared to non-edited Fielder line was much more substantial (up to 47-fold) than that reported for wild-type and edited Bobwhite (2.6-fold). This difference is mainly associated with the introgression of 1R chromosomal segment from rye into chromosome 1B of cultivar Bobwhite. This introgression resulted in the loss of gliadin- and glutenin-encoding genes located on chromosome arm 1BS (Supplementary Table 7), leading to lower immunoreactivity of cv. Bobwhite compared to cv. Fielder. The level of immunoreactivity reduction to G12 and R5 mAbs observed in our study was proportional to the number of immunoreactive peptides lost due to editing ω-and γ-gliadin gene clusters in cv. Fielder, suggesting that these mAbs primarily recognize epitopes located in these two gliadin gene families. The edited gliadin gene clusters accounted for ~30% of total gluten immunoreactivity detectable using a mix of antibodies (Lacorn et al., 2019) capable of detecting most gluten proteins.

Development of hypoimmunoreactive varieties using conventional breeding is challenged by the lack of natural gluten gene loci in wheat and its wild relatives that carry only low-toxicity epitopes. The gene editing technologies combined with new genomic resources provide unique opportunities for targeted modification of gluten genes enriched for immunoreactive peptides. The improved protein quality and reduced immunotoxicity of the edited wheat line developed here suggest that it could be directly incorporated into breeding programs as a source of new gliadin gene alleles. In addition, our study demonstrates that gene editing could be also applied to remove large gliadin gene clusters, which is an important step towards modifying natural gluten gene loci by replacing high-toxicity with low-toxicity alleles. In coming years, we could expect significant advances towards developing new wheat lines with engineered gluten gene clusters that have lower risk to human health and characterized by good end-use quality.

## Materials and methods

### Identification of gliadin genes in the genome of cultivar Fielder

A total of 17 ω-, 361 α/β- and 261 γ-gliadin genes cloned from *T. urartu* (2n=14, AA), *T. monococcum* (2n=14, AA), *Aegilops tauschii* (2n=14, DD), *T. turgidum* (2n=28, AABB) and *T. aestivum* (2n=42, AABBDD) were downloaded from the NCBI database. These 639 sequences were compared with the Fielder genome (Sato et al., 2021) using the BLASTN program. In addition, we extrapolated gene models from the RefSeq v.2.1 of cv. Chinese Spring assembly using the Liftoff program (Shumate and Salzberg, 2021). For ω-gliadin genes, this analysis identified five, eight and five gene copies on chromosomes 1A, 1B and 1D, respectively. The translated ω-gliadin amino acid sequences showed that cv. Fielder has two functional and three pseudogene copies on chromosome 1A, two functional copies and six pseudogenes on chromosome 1B, and five functional copies on chromosome 1D. For α/β-gliadin genes, there were, respectively, 11, 27 and 14 copies on chromosomes 6A, 6B and 6D. The translated amino acid sequences suggest that 10 of 11 copies, 15 of 27 copies and nine of 14 copies are functional in chromosomes 6A, 6B and 6D, respectively. For γ-gliadin genes, we detected seven, seven and nine copies on chromosomes 1A, 1B and 1D, respectively. The translated amino acid sequences suggest that five of seven copies are functional on both chromosomes 1A and 1B, and eight of nine copies are functional on chromosome 1D (Supplementary Table 6).

### CRISPR-Cas9 mediated ω- and γ-gliadin gene editing in Fielder

The sequence of nine functional ω-gliadin genes were analyzed using the web-tool CRISPOR (http://crispor.tefor.net/) (Concordet et al., 2018). The gRNAs targeting the repetitive regions within both the individual genes and across different gene copies were selected for the synthesis. Each gRNA was synthesized as two complementary oligonucleotides with four nucleotide overhangs at both termini by Integrated DNA Technologies (IDT, Coralville, IA, USA). The oligonucleotides were annealed and subcloned into the Csy4-spCas9 vector plasmid (pCas9Pvcsg) using the Golden Gate reaction with restriction endonuclease BsaI-HF v2 (New England BioLabs, Ipswich, MA, catalog #R3733L). The expression of Csy4, gRNA and spCas9 in plasmid pCas9Pvcsg is respectively driven by 35S CaMV promoter from Cauliflower Mosaic Virus, PvUbi1 promoter from switchgrass (*Panicum virgatum* L.) and ubiquitin promoter from maize (*Zea mays*). The insert of each gRNA fragment was validated by Sanger Sequencing using BigDye^™^ Terminator v3.1 Cycle Sequencing Kit (ThermoFisher Scientific, catalog #4337455).

### Biolistic transformation, plant growth and screening of mutants for the fragment deletions

Seven gRNA constructs were pooled at equimolar proportions and co-transformed with *bar-gene* vector into the immature embryos of Fielder through biolistic transformation (Saintenac et al., 2013). Genomic DNA (gDNA) was isolated from the young leaves of transgenic plants to test for the presence of Cas9 and gRNA fragments, as well as identifying the mutants with targeted fragment deletions. Briefly, leaf tissues were collected and homogenized in 400 μL of TPS buffer (100 mM Tris-HCl, 10 mM EDTA, 1 M KCl, pH8.0) using TissueLyser II (Qiagen), followed by incubation for 20 min at 75°C. After centrifugation, 130 μL of the supernatant was mixed with 130 μL of 100% isopropanol and incubated for 30 min at room temperature. DNA was precipitated, rinsed with 70%ethanol and dissolved in 150 μL of double distilled water (ddH2O). Plants were grown in 1 L square pot filled with three parts of soil mixture including soil, peat moss, perlites and CaSO4 at a volume ratio of 20:20:10:1, covered by one part of SunGro soil, and arranged according to the completely randomized design in the greenhouse of Kansas State University under 18 hr light and 6 hr dark with day and night temperature set to 25°C and 20°C, respectively (Wang et al., 2018a).

The presence of Cas9 and gRNA fragments in T_0_ plants was validated by PCR amplification using primers listed in Supplementary Table 3. The primer pair 1A1DmiseqF-1&1A1DmiseqR-4 are designed to amplify 791-bp fragment from FD1A_omega1, 872-bp fragment from FD1A_omega3, 830-bp fragment from FD1D_omega1, and 854-bp fragment from other four gene copies in chromosome 1D. The amplified fragments include two targeted regions in FD1A_omega1 and FD1A_omega3, seven regions in FD1D_omega1, and six regions in other four gene copies of chromosome 1D. The primer pairs FD1B114miseqF-3&FD1B1216miseqR-1 and FD1B114miseqF-3&FD1B1216miseqR-2 are designed to amplify 1303-bp fragments from FD1B_omega3 and 1322-bp from FD1B_omega5, respectively (Supplementary Table 4).

The PCR reaction was performed using Taq DNA Polymerase (Bullseye Taq DNA Polymerase, 1,000 Units, MidSci, SKU number: BETAQ-1000) with a reaction volume of 15 ul including 1.5 ul 10× Taq buffer, 0.8 ul 2mM dNTP, 0.9 ul 25mM MgCl2, 0.2 ul Taq polymerase, 1 ul 5pmol primer mix, 1 ul gDNA template and 9.6 ul ddH2O. The dropdown program was used for PCR setting including the initial denaturation at 94°C for 4 min, 5 cycles of 94°C for 15 s, 65°C for 30 s and 72°C for 20 s, 5 cycles of 94°C for 15 s, 60°C for 30 s and 72°C for 20 s, followed by 20 cycles of 94°C for 15 s, 55°C for 30 s and 72°C for 20 s, ended with 72°C for 5 min. The amplified products were checked by running the 2% agarose gel. The fragment deletion events of T_1_ plants were examined by PCR amplification using the primers listed in Supplementary Table 3. The PCR reaction was performed using NEBNext^®^ High-Fidelity 2× PCR Master Mix (New England BioLabs, catalog #M0541) with a reaction volume of 10 ul including 5 ul 2× PCR Master Mix, 1 ul 5pmol primer mix, 1 ul DNA template and 3 ul ddH2O. The dropdown program was used for PCR setting including the initial denaturation at 98°C for 2 min, 5 cycles of 98°C for 10 s, 65°C for 30 s and 72°C for 40 s or 1 min, 5 cycles of 98°C for 10 s, 60°C for 30 s and 72°C for 40 s or 1 min, 20 cycles of 98°C for 10 s, 55°C for 30 s and 72°C for 40 s or 1 min, ended with 72°C for 5 min. The amplified products were checked by running the 1.5% agarose gel.

### Next generation sequencing of PCR amplicons

Next generation sequencing (NGS) of PCR amplicons was also used for detecting the gene editing events in the ω-gliadin genes (Wang et al., 2018b). The genomic regions harboring the targeted regions were amplified using primer pairs 1A1DmiseqF-1&1A1DmiseqR-1 and 1BmiseqF-2&1BmiseqR-2 carrying the tails for the second round of PCR. The second round of PCR was used to finalize the reconstruction of Illumina Truseq adaptors and add barcodes for multiplexed NGS analysis of multiple genomic regions (Supplementary Table 3). All reads passing quality control were aligned against their corresponding reference sequence from wild-type Fielder genome. The editing events were visualized using Unipro UGENE v38.1 (Okonechnikov et al., 2012).

### Characterization of a gene-edited line by whole genome sequencing

The 2 x 150-bp Illumina reads have been mapped to the Fielder genome using BWA-MEM with the default settings. Read alignments have been filtered to create two alignment files: 1) with the uniquely and 2) multiply mapped reads with one mismatch per alignment allowed in both datasets. Further, the depth of read coverage was calculated using samtools. The variant calling procedure followed the best practices suggested for the GATK v.4.0 software with the default settings and was performed using the alignment file with the uniquely mapped reads.

Before analyzing gene editing events, we have identified potential sites targeted by gRNAs within the gliadin-encoding genes. For this purpose, the target sites for the designed gRNAs were located within the Fielder genome using “seqkit locate” command with up to 3 mismatches allowed. The gRNA target sites were classified into two groups: high confidence (HC) and low confidence (LC). The HC group included target sites that had no mismatches with the gRNAs or had two or fewer mismatches within a region more than 10-bp away from the PAM. The LC group included sites not included into high confidence set.

We expected that the presence of multiple gRNA target sites designed to the repetitive motifs within the gliadin genes would result in small- and large-scale deletions either with the gene cluster or within individual genes. These deletions could be detected either based on the drop of read coverage at the gRNA biding sites if editing affected only small portion of the gene or based on the drop of read coverage within the coding region if editing resulted in deletion of an entire gene or substantial part of the gene. To detect edited genes with abnormally low read coverage, we have used mean depth of read coverage per gene calculated for each gene in the Fielder genome to build null distribution. The genes below the 5^th^ percentile of mean read coverage were considered tentatively edited. We have also defined 5^th^ percentile threshold for depth of read coverage within the gRNA binding sites using the distribution build using the coverage data calculated for 21-bp long windows selected randomly within the coding regions. These thresholds were applied to read coverage data calculated for gliadin coding regions and for gRNA target sites within the gliadin genes.

In addition, we have analyzed aligned reads for the presence of small-scale deletions in the read alignments near the PAM at the gRNA biding sites. To validate these deletions, we have also re-assembled reads aligned to the gliadin genes and aligned assembled contigs to the Fielder genome. This approach allowed us to detect gene edits even in the cases when no significant drop in read coverage was observed. By combining information from GATK, indel analysis and depth of read coverage, we have identified gene editing events within the gliadin gene clusters.

We have also analyzed line 387-3-6 for the presence of off-target gene editing events. For this analysis we have used sites outside of the gliadin genes aligning to gRNAs with 1, 2 or 3 mismatches. Then we checked for overlap of these sites with the indels detected by GATK.

### Extraction of gliadins, SDS-extractable and SDS-unextractable polymeric proteins

Gliadins were isolated from the half seeds with endosperm from the edited lines and wild-type Fielder. The half seeds were ground into wholemeal flour using TissueLyser II, followed by adding 1 ml of 70% ethanol (Yu et al., 2021). After shaking at room temperature for 1 hr and centrifuging at 13,000 rpm for 15 min, the supernatant containing soluble gliadin fraction was collected.

SDS-extractable and SDS-unextractable polymeric proteins were also extracted from the half seeds with endosperm. Two half seeds were ground together using TissueLyser II to obtain 30 mg wholemeal flour for each sample, followed by adding 1.5 ml of 0.05 M phosphate buffered saline buffer (PBS, pH 6.9) with 0.05% sodium dodecyl sulphate (SDS, Sigma-Aldrich). After shaking at room temperature for 1 hr, the supernatant was collected as SDS-extractable polymeric protein (EPP) fraction by centrifuging at 13,000 rpm for 15 min. Another 0.6 ml of 0.05 M PBS extraction buffer with 0.05% SDS was added to the pellets for isolating SDS-unextractable polymeric proteins (UPP). The pellets were suspended in the extraction buffer, followed by sonication using a probe sonication instrument at 10 W for 3 min. After shaking for 30 min, the supernatant was collected as UPP by centrifuge at 13,000 rpm for 15 min (Batey et al., 1991).

### Characterization of gliadins, SDS-extractable and SDS-unextractable polymeric proteins using high performance liquid chromatograph

The characterization of gliadins was performed by reverse-phase high performance liquid chromatograph (RP-HPLC) using an Agilent 1290 Infinity II LC system (Agilent Technologies, http://www.agilent.com). Twenty microliters of the gliadin extracts were injected into a C18 reversed-phase Zorbax 300 StableBond column (4.6×250 mm, 5 μm, 300 Å, Agilent Technologies), maintained at 60°C. The eluents were ultrapure water (solvent A) and acetonitrile (ACN, LC-MS Grade, Thermo Scientific, Catalog #A955-4, solvent B), each containing 0.06% Trifluoroacetic Acid (TFA, LC-MS Grade, Thermo Scientific, Catalog #85183). The flow rate was set at 1 ml min^-1^. Gliadins was characterized by a linear gradient from 21% to 48% of solvent B in 55 min and detected by UV absorbance at 214 nm. After each run, the column was balanced for 15 min.

The characterization of EPP and UPP were performed by size-exclusion high performance liquid chromatograph (SE-HPLC) using an Agilent 1290 Infinity II LC system (Agilent Technologies, http://www.agilent.com). Twenty microliters of the EPP and UPP extracts were separately injected into a Bio SEC-5 column (4.6×300 mm, 500 Å, Agilent Technologies), maintained at 25°C. The eluents were ultrapure water (solvent A) and ACN (LC-MS grade, solvent B), each containing 0.1% TFA (LC-MS grade). The flow rate was set at 0.35 ml min^-1^. The EPP and UPP were characterized by using a constant gradient with 50% of solvent A and 50% of solvent B in 20 min, and detected by UV absorbance at 214 nm.

### Quantification of gliadin subtypes, components of SDS-extractable and SDS-unextractable polymeric proteins

Chromatograms were managed by OpenLAB CDS ChemStation Edition for LC & LC/MS Systems v2.18.18 (Agilent Technologies). Based on hydrophobicity, three obvious boundary areas corresponding to ω-gliadins, α/β-gliadins and γ-gliadins could be identified in the gliadin chromatogram. There were three separated areas within ω-gliadins, sequentially assigned as ω5-gliadins, albumins/globulins and ω1,2-gliadins. The amount of each gliadin subtype was calculated as the area under the chromatogram trace of each component. Thus, the percentage of ω5-, ω1,2-, ω- and γ-gliadins were calculated as the area under each component respectively divided by the total area of gliadins without the area under albumins/globulins.

The peaks in the chromatograms of both SDS-extractable and SDS-unextractable polymeric proteins corresponding to glutenins, gliadins, and albumins/globulins are shown in Fig. 3c. Calculation of the amount of each component was performed based on the area-under-the-peak method. Thus, the ratio of glutenins to gliadins was calculated as the glutenins in both EPP and UPP divided by the gliadins in both EPP and UPP; the ratio of polymeric to monomeric proteins was calculated as the glutenins in both EPP and UPP divided by the gliadins and albumins/globulins in both EPP and UPP; the percentage of GMP was calculated by the glutenins in UPP divided by the glutenins in both EPP and UPP.

### Enzyme-Linked Immunosorbent Assay

The Enzyme-Linked Immunosorbent Assay (ELISA) was conducted using the G12 mAb kit (AgraQuant^®^Gluten G12^®^, 10001994, Romer Labs, Newark DE, USA), the R5 mAb kit (RIDASCREEN^®^Gliadin, Art. No. R7001, R-Biopharm, Darmstadt, Germany) and the R-BioPharm Total Gluten kit (RIDASCREEN^®^Art. No. R7401). This assay contains a mixture of four mAbs: mAb R5 as well as mAbs to the HMW-GS, LMW GSs and rye seaclins (Lacorn et al., 2019). The sample preparation and measurements were conducted following the manufacturer’s instructions for the kits.

### Comparison of wheat 1BS and rye 1RS

A comparison in the numbers of toxic epitopes binding to R5 and G12 mAbs between gliadins in wheat 1BS and secalins in rye 1RS was made, shown as 132 vs 95 for 1BS vs 1RS in R5 and 54 vs 59 for 1BS vs 1RS in G12. Moreover, there was no orthologs of low molecular weight glutenin subunit (LMW-GS) found in 1RS (Li et al., 2021). The LMW-GS encoding genes are a cluster of genes which are tightly linked with gliadin gene loci in short arm of chromosome 1A, 1B and 1D. The sequence similarity between LMW-GS and gliadin gene is mainly evident in the parts that encode the repetitive domains of protein, which is the toxic epitope composed of PxQ motifs (D’Ovidio and Masci, 2004). Therefore, in comparison with Fielder, the wild-type Bobwhite with 1BL/1RS translocation used in Sánchez-León et al. 2018 is already a low gluten toxicity cultivar.

### Statistical analysis

The boxplot was used to visualize the differences in the investigated parameters between wild-type Fielder and gene edited lines. The mean value, interquartile ranges (IQR), minimal value and the maximum value were indicated in the boxplots. The end of the top line and the bottom line of the box were the third quartile (Q3) + 1.5× IQR and the first quartile (Q1) – 1.5× IQR, respectively. The two-sided *t*-test was carried out to assess the statistical significance of the differences in the investigated parameters between wild-type Fielder and the gene edited lines.

## Author Contributions

ZY – designed and conducted gene editing experiments, identified gene-edited plants, designed and conducted experiments to characterize flour quality using HPLC, contributed to drafting the manuscript; UY –analysis of gene editing events using whole genome sequencing data; MT – conducted analyses of the immunoreactivity of the flour protein extracts; AF – contributed resources and tools for the HPLC analysis of flour quality traits; AA – generation of amplicon and whole genome next-generation sequencing data for detecting gene editing events; HT – generation of transgenic plants for gene editing; EA – conceived idea, designed gene editing experiments, analyzed and interpreted results, supervised the project and wrote the manuscript.

## Acknowledgement

This project was supported by the Agriculture and Food Research Initiative Competitive Grants 2021-67013-34174 and 2020-67013-30906 from the USDA National Institute of Food and Agriculture, and grants from the Bill and Melinda Gates Foundation (INV-004430) and Kansas Wheat Commission. We thank Hyeonju Lee for conducting plant transformation and regeneration, Shuangye Wu for preparing the genomic library of edited cultivar Fielder, Guihua Bai for providing access to Sanger sequencing instrumentation and Dwight Davidson for help with greenhouse management. Mention of trade names or commercial products in this publication is solely for the purpose of providing specific information and does not imply recommendation or endorsement by the U.S. Department of Agriculture. USDA is an equal opportunity provider and employer.

## Conflict of interest

The authors declared that they do not have conflict of interests.

## Supporting information

Supplementary Table 1. Number of R5 and G12 mAbs binding toxic epitopes detected within the gliadin-encoding genes from four wheat cultivars including Fielder, Kariega, Chinese Spring and LongReach Lancer.

Supplementary Table 2. Potential gRNA target sites within the gliadin genes.

Supplementary Table 3. List of PCR primers used in the study.

Supplementary Table 4. Primers for PCR-based screening of fragment deletions.

Supplementary Table 5. NGS based detection for editing events by PCR amplicon sequencing.

Supplementary Table 6. Summary of gene editing events detected by whole genome sequencing in the ω- and γ-gliadin gene clusters.

Supplementary Table 7. The differences in the functional storage protein gene copy number between wheat 1BS and Rye 1RS.

Supplementary Table 8. Raw data for generating figures to show the impacts of gene editing on the content of each gliadin subtype, parameters of protein extracts correlated with grain protein quality for breadmaking and immunoreactivity in Figure 3.

## Notes

### Competing Interest Statement

The authors have declared no competing interest.

