## Supplementary Information for "CRISPR-based editing of the ω- and γ-gliadin gene clusters reduces wheat immunoreactivity without affecting grain protein quality"

### Supplementary Figures

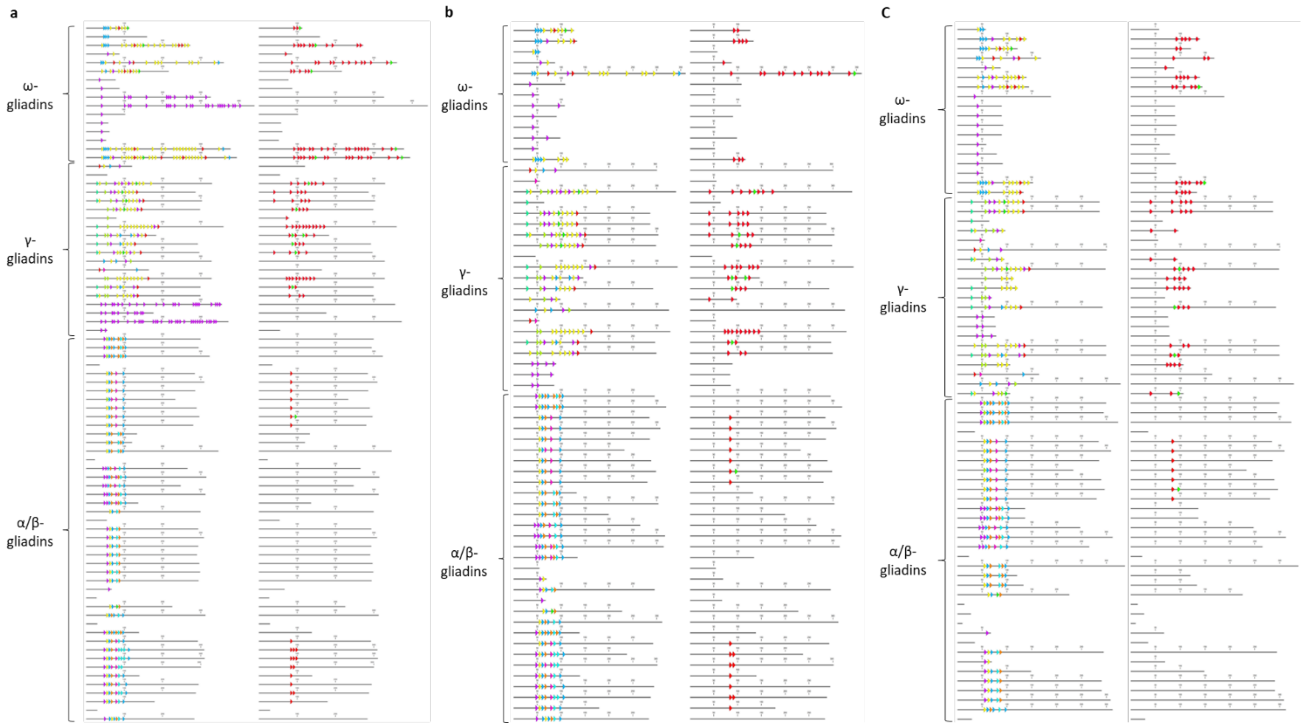


**Supplementary Figure 1.** The distributions of the 17 toxic epitopes including 11 binding to R5 mAb and six binding to G12 mAb across all three gliadin subtypes of cultivars Kariega, Chinese Spring and LongReach Lancer. The 17 toxic epitopes were indicated by the arrows with corresponding color shown in Fig. 1a. The left and right graphs in the panels **a, b** and **c** were, respectively, referred to the distributions of the toxic epitopes binding to R5 and G12 mAbs for cv. Karriega (**a**), cv. Chinese Spring (**b**), and cv. LongReach Lancer (**c**). The number of toxic epitopes binding to R5 and G12 mAbs across all three gliadin subtypes of each cultivar are shown in Supplementary Table 1.

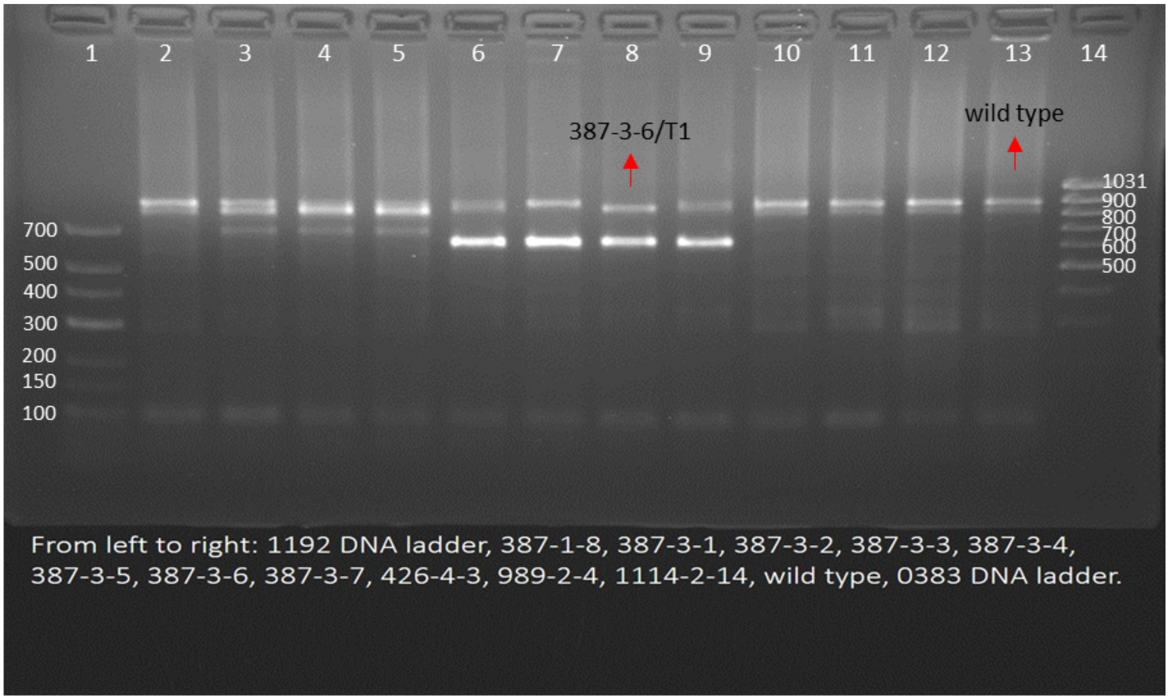


**Supplementary Figure 2.** PCR-based screening of fragment deletions in five functional ω-gliadin gene copies from chromosomes 1A and 1D in the $T_1$ population using primer pair 1A1DmiseqF-1&1A1DmiseqR-4. lane 1: 1192 DNA ladder; lane 2: 387-1-8; lane 3: 387-3-1; lane 4: 387-3-2; lane 5: 387-3-3; lane 6: 387-3-4; lane 7: 387-3-5; lane 8: 387-3-6; lane 9: 387-3-7; lane 10: 426-4-3; lane 11: 989-2-4; lane 12: 1114-2-14; lane 13: wild-type Fielder; lane 14: 0383 DNA ladder; the negative control was wild-type Fielder in lane 13, indicated by red arrow. The expected size of PCR products for the control are 791-bp fragment from FD1A_omega1, 872-bp fragment from FD1A_omega3, 830-bp fragment from FD1D_omega1, and 854-bp fragment from other four gene copies in chromosome 1D. The amplified products from seven $T_1$ plants derived from $T_0$ line 387-3 including plants 387-3-1, 387-3-2, 387-3-3, 387-3-4, 387-3-5, 387-3-6, 387-3-7 had fragments with smaller size than the control, indicating there were fragment deletions in the ω-gliadin gene copies located on chromosomes 1A and 1D of theses seven $T_1$ plants. Among them, the amplified fragments from $T_1$ plant 387-3-6 in lane 8 had two bands, which were both smaller than the control, indicating that both were amplified from the edited gene copies.

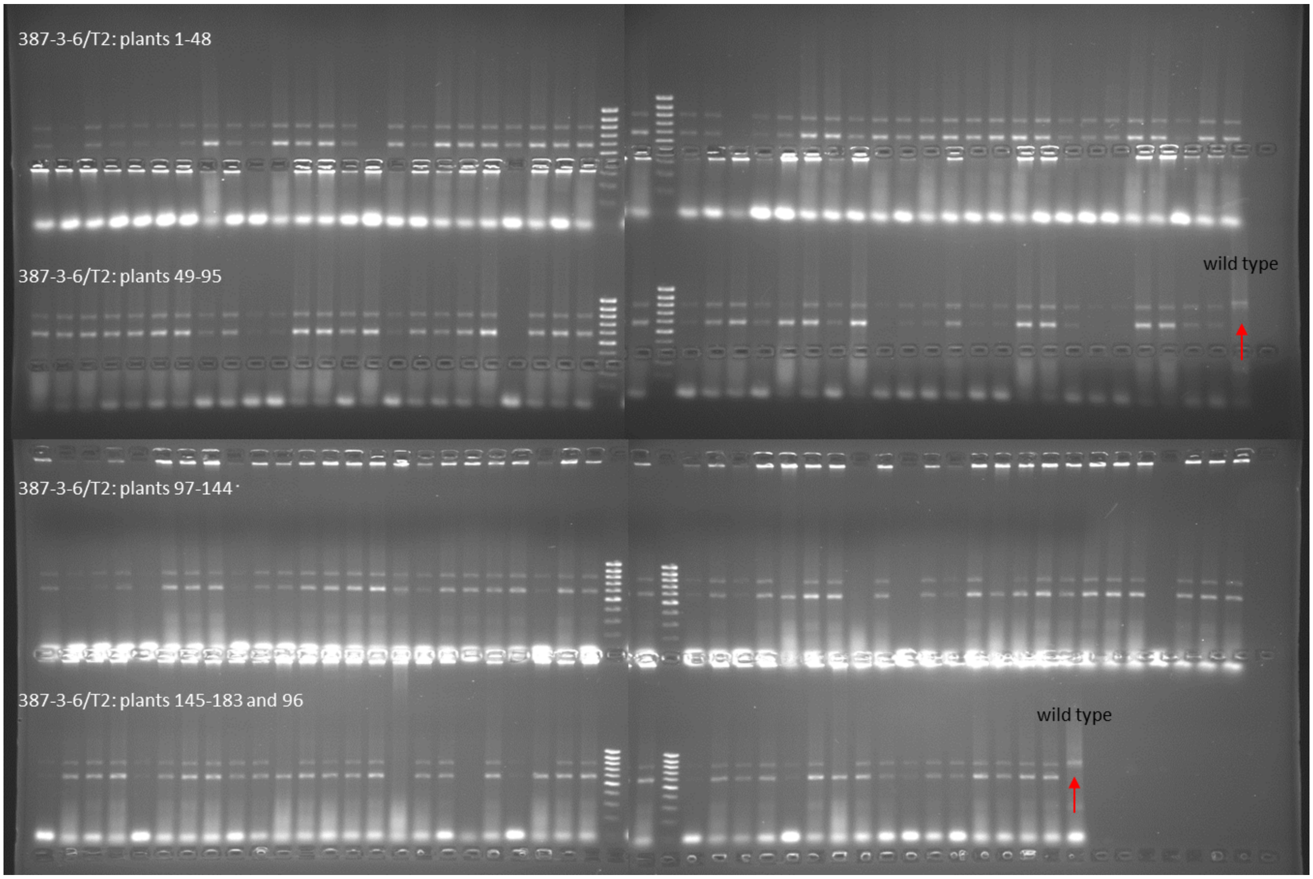


**Supplementary Figure 3.** PCR-based screening of fragment deletions in five functional ω-gliadin gene copies located on chromosomes 1A and 1D in the $T_2$ population including 183 lines derived from plant 387-3-6. PCR was performed using primer pair 1A1DmiseqF-1&1A1DmiseqR-4. The 0383 DNA ladder was loaded into the twenty-fifth lane of each row. The negative control was wild-type Fielder, which is marked by the red arrow. The amplified fragments from $T_2$ plants 1-48, 49-95, 97-144, 145-183 and 96 were shown in the four rows from top to bottom. The amplified fragments from each $T_2$ plant had two bands of smaller size than the control, indicating both were amplified from the edited gene copies. There was no segregation in the fragment deletions observed among the analyzed 183 lines, revealing that the $T_1$ line 387-3-6 is homozygous for the editing events.

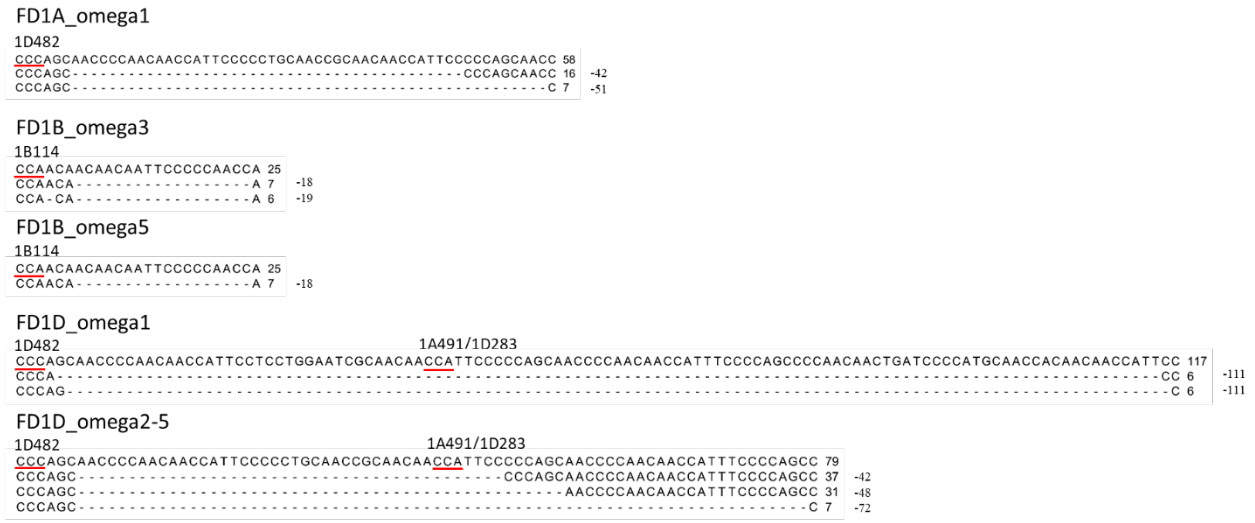


**Supplementary Figure 4.** NGS-based detection for fragment deletions in the ω-gliadin genes of edited line 387-3-6 using PCR amplicons produced by primer pairs 1A1DmiseqF-1&1A1DmiseqR-1 and 1BmiseqF-2&1BmiseqR-2. The PAM site NGG is underlined. The deleted bases were shown by black dots. The sizes of the deleted fragments are shown as negative number at the end of sequence. The sizes of the deleted fragments in FD1A_omega3 were not detected within a total of 17 reads using the PCR amplicons based NGS, mostly due to the occurrence of the long fragment deletion resulting in the targeted amplified region being deleted (Fig. 2, Supplementary Table 6). A summary of the editing events detected by PCR amplicons based NGS in each copy was shown in Supplementary Table 5.

### Supplementary Tables

**Supplementary Table 1.** Number of R5 and G12 mAbs binding toxic epitopes detected within the gliadin-encoding genes from different wheat cultivars.

| Sequence | Cultivar | Gliadin subtype | R5 | G12 |
|---|---|---|---|---|
| Complete sequence | Fielder | ω-gliadins | 253 | 128 |
| | | α/β-gliadins | 263 | 26 |
| | | γ-gliadins | 234 | 92 |
| | Kariega | ω-gliadins | 163 | 72 |
| | | α/β-gliadins | 245 | 22 |
| | | γ-gliadins | 195 | 64 |
| Incomplete sequence | Chinese Spring | ω-gliadins | 54 | 23 |
| | | α/β-gliadins | 185 | 18 |
| | | γ-gliadins | 140 | 61 |
| | LongReach Lancer | ω-gliadins | 85 | 33 |
| | | α/β-gliadins | 156 | 8 |
| | | γ-gliadins | 113 | 45 |

**Supplementary Table 2.** Potential gRNA target sites within the gliadin genes (external Excel file).

**Supplementary Table 3.** List of PCR primers used in the study.

| Purpose | Primer | Sequence | Amplified fragment length |
|---|---|---|---|
| Fragment deletions | 1A1DmiseqF-1 | CTCTTTCCCTACACGACGCTCTTCCGATCTCGCTTCCCAGACCCAACAATCGT | 791, 830, 854, 872-bp |
| | 1A1DmiseqR-4 | CTGGAGTTCAGACGTGTGCTCTTCCGATCTGCTTACCACCGATGCTTGTAAGACTA | |
| | FD1B114miseqF-3 | CTCTTTCCCTACACGACGCTCTTCCGATCTCGCTTGGAATTGCATACTCCACAAGA | 1303-bp |
| | FD1B1216miseqR-1 | CTGGAGTTCAGACGTGTGCTCTTCCGATCTGCTTAAGGCCACTGATACTTATAACGT | |
| | FD1B114miseqF-3 | CTCTTTCCCTACACGACGCTCTTCCGATCTCGCTTGGAATTGCATACTCCACAAGA | 1322-bp |
| | FD1B1216miseqR-2 | CTGGAGTTCAGACGTGTGCTCTTCCGATCTGCTTATCATAGGCCACAGATACTTA | |
| Cas9 fragment | SpCas9F2 | agattacgaaggctccgctc | 296-bp |
| | SpCas9R2 | gctgccgttatcgaatgtcc | |
| gRNA fragment | Pvintron1F1 | TACCAATGATGACCTTATCTCTC | 198-bp |
| | sgScaffoldR1 | TTCAAGTTGATAACGGACTAGC | |
| NGS | 1A1DmiseqF-1 | CTCTTTCCCTACACGACGCTCTTCCGATCTCTGTACCCAGACCCAACAATCGT | 301, 325-bp |
| | 1A1DmiseqR-1 | CTGGAGTTCAGACGTGTGCTCTTCCGATCTGCTTAGGTTGTTGGAGTTCAGGAAATAA | |
| | 1BmiseqF-2 | CTCTTTCCCTACACGACGCTCTTCCGATCTCTGTACAAGGAATTGCATACTCCACAA | 306-bp |
| | 1BmiseqR-2 | CTGGAGTTCAGACGTGTGCTCTTCCGATCTGCTTAAATTCCTGTTGCGGCAATT | |
| | PCR_Truseq_Amp_F | AATGATACGGCGACCACCGAGATCTACACTCTTTCCCTACACGAC | |
| | PCR_Truseq_Amp_R_21 | CAAGCAGAAGACGGCATACGAGATCGAAACGTGACTGGAGTTCAGACG | |
| | PCR_Truseq_Amp_R_23 | CAAGCAGAAGACGGCATACGAGATCCACTCGTGACTGGAGTTCAGACG | |

**Supplementary Table 4.** Primers for PCR-based screening of fragment deletions.

| Primer | Gene | Chromosome | Amplified fragment length | Number of targeted region |
|---|---|---|---|---|
| 1A1DmiseqF-1&1A1DmiseqR-4 | FD1A_omega1 | 1A | 791-bp | 2 |
| | FD1A_omega3 | 1A | 872-bp | 2 |
| | FD1D_omega1 | 1D | 830-bp | 7 |
| | FD1D_omega2 | 1D | 854-bp | 6 |
| | FD1D_omega3 | 1D | 854-bp | 6 |
| | FD1D_omega4 | 1D | 854-bp | 6 |
| | FD1D_omega5 | 1D | 854-bp | 6 |
| FD1B114miseqF-3&FD1B1216miseqR-1 | FD1B_omega3 | 1B | 1303-bp | 4 |
| FD1B114miseqF-3&FD1B1216miseqR-2 | FD1B_omega5 | 1B | 1322-bp | 4 |

**Supplementary Table 5.** NGS based detection of editing events by sequencing PCR amplicons.

| Primer | Gene | Chromosome | Number of targeted regions | Type of editing event | Size of event |
|---|---|---|---|---|---|
| 1A1DmiseqF-1&1A1DmiseqR-1 | FD1A_omega1 | 1A | 2 | Deletion | 42- and 51-bp |
| | FD1A_omega3 | 1A | 1 | Deletion | N/A* |
| | FD1D_omega1 | 1D | 3 | Deletion | 111-bp |
| | FD1D_omega2 | 1D | 3 | Deletion | 42-, 48-, and 72-bp |
| | FD1D_omega3 | 1D | 3 | Deletion | 42-, 48-, and 72-bp |
| | FD1D_omega4 | 1D | 3 | Deletion | 42-, 48-, and 72-bp |
| | FD1D_omega5 | 1D | 3 | Deletion | 42-, 48-, and 72-bp |
| 1BmiseqF-2&1BmiseqR-2 | FD1B_omega3 | 1B | 1 | Deletion | 18- and 19-bp |
| | FD1B_omega5 | 1B | 1 | Deletion | 18-bp |

N/A*: The size of the deleted fragments in FD1A_omega3 were not detected within a total of 17 reads using the PCR amplicons based NGS, mostly due to the occurrence of the long fragment deletion resulting in the targeted amplified region being deleted (Fig. 2, Supplementary Table 6).

**Supplementary Table 6.** Summary of gene editing events detected by whole genome sequencing in the ω- and γ-gliadin gene clusters (external Excel file).

**Supplementary Table 7.** The differences in the functional storage protein gene copy number between wheat 1BS and rye 1RS.

| Species | cultivar | ω-gliadins/secalins | γ-gliadins/secalins | LMW-GS |
|---|---|---|---|---|
| Wheat 1BS | Fielder | 2 | 8 | 4 |
| Rye 1RS | Weining | 3 | 5 | 0 |

**Supplementary Table 8.** Raw data for generating figures to show the impact of gene editing on the content of each gliadin subtype, parameters of protein extracts correlated with grain protein quality for breadmaking and immunoreactivity in Figure 3 (external Excel file).
